# Detection dog performance state estimation from pre-stimulus video and physiological signals using deep learning and Bayesian inference

**DOI:** 10.1101/2025.10.31.685787

**Authors:** Jörg Schultz, Liza Rothkoff, Edgar Aviles-Rosa, Nathaniel J. Hall, Michele N. Maughan

## Abstract

Detection dogs play a critical role in operational settings ranging from explosives detection to medical diagnostics. Their unmatched olfactory capabilities allow them to locate trace-level targets that remain beyond the reach of current technology. However, detection performance can decline without overt behavioral signs, posing a challenge for timely and informed deployment decisions. Identifying subtle precursors to missed alerts remains an open problem with both practical and computational significance.

To address this, we recorded detection dogs on a treadmill-based scent delivery system under tightly controlled conditions, enabling precise timing of odor presentation. We focused on the 4.5 seconds preceding each odor pulse to capture anticipatory movement through video and physiological state via sensors. Using markerless pose estimation, we extracted high-resolution joint trajectories, while concurrent recordings provided heart rate, heart rate variability, and core temperature. We trained a spatiotemporal deep learning model that integrates Graph Attention Networks and Temporal Convolutional Networks to predict missed indications from these short, pre-stimulus windows. The model successfully identified 85% of missed alerts with 81% precision. Heart rate variability emerged as the most informative physiological input, suggesting a strong autonomic component to declining readiness.

To move from isolated predictions to actionable insights, we implemented a Bayesian aggregation framework that estimates each dog’s latent miss rate over time. This probabilistic formulation enables adaptation to individual baselines and operational risk thresholds, supporting context-sensitive deployment decisions.

While data were collected in a laboratory setting, our findings highlight behavioral and physiological signatures that precede performance failure. This work lays the foundation for real-time readiness monitoring systems that integrate wearable sensing with interpretable machine learning, supporting timely and welfare-conscious deployment decisions.

**Author summary:** Detection dogs are used in high-stakes situations—from finding explosives to diagnosing diseases. But detection work is physically and mentally demanding, and a dog’s performance can quietly decline over time. One challenge is that handlers often can’t tell when a dog is starting to lose focus or miss targets until it’s too late. In our study, we used a treadmill-based setup to record dogs as they performed scent detection tasks. We tracked how they moved, monitored their heart rate and temperature, and trained a computer model to recognize patterns that typically show up just before a missed target. Importantly, we built a second layer that transforms the model’s output into a running estimate of how likely the dog is to miss a target—based on its recent behavior. This second layer can be tuned to the task and the dog. It can issue earlier warnings during high-risk missions like explosives detection and delay warnings for experienced dogs known to perform reliably even under physical and mental strain. The result is a flexible framework—a first step toward tools that could support better decision-making in real-world deployments.

## Introduction

Detection dogs are indispensable assets in diverse operational domains, including explosives (1) and drugs (2) interdiction, search-and-rescue (3), wildlife conservation (4), and medical detection (5). Their extraordinary olfactory capabilities allow them to locate targets that remain beyond the reach of technological sensors (6). In many of these scenarios, reliability is critical: a single missed alert can compromise mission success, endanger human lives, or undermine confidence in a canine program. Even for highly trained and physically fit dogs, detection performance can vary across and within working sessions, influenced by factors such as physiological state (7; 8), environmental conditions (9), and task demands. The consequences of performance decline make it essential for handlers to detect early signs of reduced readiness and to act before an operational failure occurs.

In the field, handlers monitor their dogs closely, looking for subtle cues in movement, posture, or search behavior that might signal fatigue, distraction, or waning motivation. This visual assessment is a cornerstone of operational decision-making, drawing on the handler’s experience and knowledge of the individual dog. However, in complex or high-pressure environments, these cues can be fleeting, occur outside the handler’s line of sight, or be overshadowed by competing demands such as navigating the search area or communicating with a team. Moreover, performance decline can begin at physiological or sensory stages that are not easily observable by the handler.

Experimental work has shown that exercise can impair olfactory detection limits before overt behavioral signs emerge, particularly at low target odor concentrations (10).

These conditions are common in operational scenarios, where targets may emit only trace quantities of volatile compounds, increasing the likelihood that early declines in detection performance go unnoticed until a missed alert occurs. This motivates the search for objective, sensor-based approaches that can complement handler expertise by capturing and quantifying behavioral and physiological indicators in real time.

From a computational perspective, this presents a challenging inference problem: estimating latent performance states from short, noisy time series of movement and physiological data, where labels are sparse and behavioral indicators are subtle.

Movement offers a particularly informative window into internal state, as it reflects subtle changes in posture, coordination, and engagement. Among available modalities, video data provides the temporal and spatial resolution needed to capture these behavioral dynamics objectively. Within the spectrum of computer vision techniques, markerless pose estimation has emerged as a powerful tool for analyzing canine movement (11; 12; 13; 14). This approach can quantify joint-level kinematics from ordinary video without the need for reflective markers or dedicated motion-capture laboratories, making it well suited for controlled studies where standardized, high-quality recordings are possible. In our work, markerless pose estimation enables us to describe movement patterns in fine detail, opening the door to identifying small but meaningful deviations associated with changes in performance readiness. Other modalities for quantifying movement, such as wearable inertial measurement units (IMUs) and accelerometers, have been successfully used to classify canine activities in operational environments (15; 16; 17; 18). While these were not part of the present study, they represent a promising avenue for translating laboratory-derived movement indicators into practical, field-ready tools.

Physiological measures offer a complementary perspective on readiness to detect and correctly alert to target odors. Core body temperature is a direct indicator of thermoregulatory load and can rise well before obvious behavioral signs of fatigue appear (19; 20; 21; 22). Heart rate (HR) and heart rate variability (HRV) index autonomic nervous system activity and have been linked to arousal, stress, and fatigue in dogs (23; 24; 25), although interpretation must account for concurrent movement. Elevated physiological strain can impair olfactory performance and shorten effective working time, even when the dog continues to appear behaviorally engaged (7; 8; 10; 26). By integrating movement-derived and physiological signals, it is possible to characterize both the external and internal states of the working dog, potentially identifying early-warning patterns that are not apparent through behavior or physiology alone.

Studying these relationships directly in real-life search scenarios presents substantial challenges. Operational searches involve many uncontrolled variables—environmental conditions, search area structure, target placement, and dog–handler interactions—that can obscure subtle performance-related patterns. Video analysis of freely moving dogs is technically demanding in such contexts, with occlusions, variable lighting, and changes in camera perspective reducing data quality. Furthermore, it is often impossible to determine the precise moment when a dog first encounters target odor, making any analysis of “pre-odor” behavior inherently uncertain. To address these issues, we used a controlled olfacto-treadmill setup in which the dog moves “in place” while odor stimuli are delivered in precisely timed and quantified pulses (10). This design allows us to control exercise intensity, regulate odor concentration (27), and standardize presentation timing, while simultaneously enabling high-quality video capture and synchronized physiological recording. Such a setup offers the experimental control necessary to isolate and analyze the specific behavioral and physiological signatures that precede detection failures.

In this study, our goal is to test whether it is possible to predict an impending missed indication from short windows of combined video-based movement data and physiological measurements. We address this question using a deep learning framework capable of extracting and integrating complex spatial–temporal patterns from these complementary data streams. A common critique of such neural network models is that they operate as “black boxes,” offering little transparency about which features drive their predictions (28). To counter this limitation, we apply complementary interpretability techniques to identify which aspects of movement and physiology the model relies on. This serves two purposes: (i) it allows us to assess whether the model’s focus aligns with behavioral cues that experienced handlers themselves attend to, thereby increasing confidence in its practical relevance, and (ii) it may help to home in on the underlying physiological mechanisms that precede performance decline.

For such a prediction framework to be useful in operational practice, it must also be adaptable to the individual dog and to the specific mission context. Dogs differ in their baseline physiology, working style, and resilience under sustained workload. A well-conditioned, experienced dog with a long record of successfully detecting very low odor concentrations may maintain effective performance longer than a young dog in its first deployment, whose endurance and behavioral consistency are still developing.

These differences mean that a “one-size-fits-all” decision threshold for performance decline is unlikely to be optimal.

The operational setting further shapes how predictions should be interpreted. The consequences of a missed detection vary widely between contexts, and so does the tolerance for risk. In high-risk missions—such as those involving improvised explosive devices—handlers may justifiably prefer to rotate or rest a dog at the earliest sign of possible decline, even if this causes operational delays or requires additional resources. In other settings, such as customs inspections or searches for small amounts of narcotics, cash, or other contraband, handlers might balance the risk of a missed alert differently, considering operational efficiency and resource availability alongside detection accuracy. In all cases, the decision to pause or continue a search is shaped by a combination of the dog’s current state, its track record, and the mission’s risk profile. Therefore, any predictive system should be flexible enough to accommodate these factors.

Taken together, our aim in this study is to determine whether deep learning methods applied to short windows of video-based movement data and physiological measurements can predict an impending missed indication. Beyond making single-trial predictions, we place these outputs into a broader decision-making context that reflects the dog’s evolving operational state. By framing this state in probabilistic terms, our approach can incorporate prior knowledge about the individual dog’s baseline performance and training history, as well as mission-specific tolerance for missed detections. Such a framework offers both the flexibility to adapt to diverse operational requirements and the interpretability needed for effective integration into handler decision-making. By enabling earlier rest or reassignment when subtle signs of fatigue emerge, it supports more informed and welfare-conscious deployment decisions.

## Material and methods

### Ethics Statement

The study was conducted at the Texas Tech University Canine Olfaction Research and Education (CORE) Lab and approved by the Texas Tech University Institutional Animal Care and Use Committee (IACUC) and the U.S. Army Animal Care and Use Review Office (ACURO).

### Data Collection and Preprocessing

#### Subjects

Four dogs participated in this study (Table 1). All dogs were between 2 and 4 years of age, healthy, and actively trained for detection work. One dog (Charles) had also participated in our prior treadmill study (10). Dogs were selected based on their familiarity with treadmill walking and consistent participation across sessions. This information has previously been reported in a companion manuscript (29).

**Table 1.** Demographic information of study dogs.

| Dog Name | Sex | Breed | Age<br>(years) | Weight<br>(kg) |
| --- | --- | --- | --- | --- |
| Adele | Spayed female | Belgian Malinois | 3 | 22 |
| Boomer | Intact male | English Springer Spaniel | 2 | 23 |
| Charles | Neutered male | Mixed breed | 4 | 20 |
| Iitooma | Neutered male | English Springer Spaniel | 3 | 17 |

#### Experimental Setup and Data Acquisition

##### Olfacto-treadmill setup and task design

All recordings were obtained during a treadmill-based odor discrimination task using a custom-built olfacto-treadmill, previously described in detail (10; 29). Dogs maintained a consistent gait—either walking (4 km/h) or trotting (8 km/h)—throughout the session, while odors were delivered via an automated olfactometer. All analyses in this study focused exclusively on the moderate-intensity trot condition (8 km/h).

##### Trial structure

Each trial began with computer-controlled activation of an odor valve (target or diluent). Dogs were trained to insert their muzzles into an odor port, breaking an infrared beam. Responses were scored as hits if the beam was broken for a dog-specific minimum duration (0.4 s for Boomer and Charles; 0.8 s for Adele and Iitooma). A failure to respond within 10 s was scored as a miss. Correct responses (both go and no-go) were reinforced with food. Trials were initiated every 20 s, yielding fixed onset intervals and uniform session durations. Target presence was randomized at 50% frequency. The experimenter was blind to trial type; valve selection and feedback tones were fully automated. The target odor was 1-bromooctane (1-BO), diluted in mineral oil (v/v).

##### Session formats

Two session types contributed data: (i) *Fixed-pace sessions* included 100 trials at a constant gait (walk or trot), lasting 33 min 20 s. (ii) *Pace-transition sessions* included 140 trials (approx. 46 min) with a mid-session gait switch (walk → trot or trot → walk) after trial 70. Each dog completed four fixed-pace sessions (two walk, two trot) and eight pace-transition sessions (four of each transition type) on separate days, with at least 18 h between sessions. Exercise was voluntary; if a dog stepped off the treadmill, the session resumed once the dog re-engaged.

##### Video recording

Video was recorded from a fixed lateral camera positioned at mid-body height under consistent indoor lighting and background. Recordings were time-synchronized with the olfactometer and physiological event logs. For each target-present trial, the 4.5 s immediately preceding odor onset were extracted for modeling. Details of pose estimation and feature extraction are provided in the following section.

##### Physiological measurements

Core body temperature (*Temp*) was measured using an ingestible AniPill^TM^ capsule (BodyCap, France), sampled every minute. Heart rate (HR) and inter-beat intervals (IBI) were continuously recorded using a Polar^@^ H10 chest strap and logged at 1 Hz. All physiological data were aligned to trial timing. HR data from one dog (Iitooma) were excluded due to intermittent sensor contact during trotting sessions.

##### Subset used for modeling

To isolate predictors of missed indications under moderate workload, we restricted analyses to target-present trials at 8 km/h. For each included trial, video and physiological data were aligned to the 4.5 s pre-odor window, and behavioral labels were assigned based on the infrared beam-break criteria described above.

#### Physiological Data Processing

Gastrointestinal temperature data were linearly interpolated to 1-second resolution and aligned to trial timestamps. Heart rate was recorded continuously at 1 Hz and similarly aligned. For modeling, each trial was represented by a 4.5-second window preceding odor onset. This window was selected to capture changes in physiology and movement prior to stimulus delivery. Trials containing signal loss or sensor dropout were excluded from analysis.

#### Trial Labeling and Ground Truth

Each trial was labeled based on the dog’s behavioral response during odor presentation. Only trials presenting a target odor (1-bromooctane) were included in the modeling analysis; diluent control trials (mineral oil) were excluded. A correct nose poke within the predefined, dog-specific duration window was labeled as a “hit,” while failure to respond was labeled as a “miss.” Labels were derived directly from the experiment control software using infrared beam breaks with individualized thresholds.

#### Data Preprocessing and Inclusion Criteria

Only trials containing a target odor with complete and valid pose and physiological data were included in the analysis. For each retained trial, we extracted a 4.5-second window immediately preceding odor onset. All input features were aligned to this window and preprocessed as follows.

#### Pose data

Pose annotations consisted of 150 frames sampled at 30 Hz, with 29 anatomical landmarks per frame (for a complete list see Figure 1). For each landmark, we extracted two pixel coordinates (x, y) specifying its location in the video frame and a likelihood value reflecting the confidence of the pose prediction. All sequences were truncated to exactly 150 frames to ensure a consistent input length across trials. To remove global positional bias, coordinates were translated such that the withers (joint 4) were located at the origin in every frame. This normalization ensured the model focused on joint movement rather than location on the treadmill.

**Fig 1.**
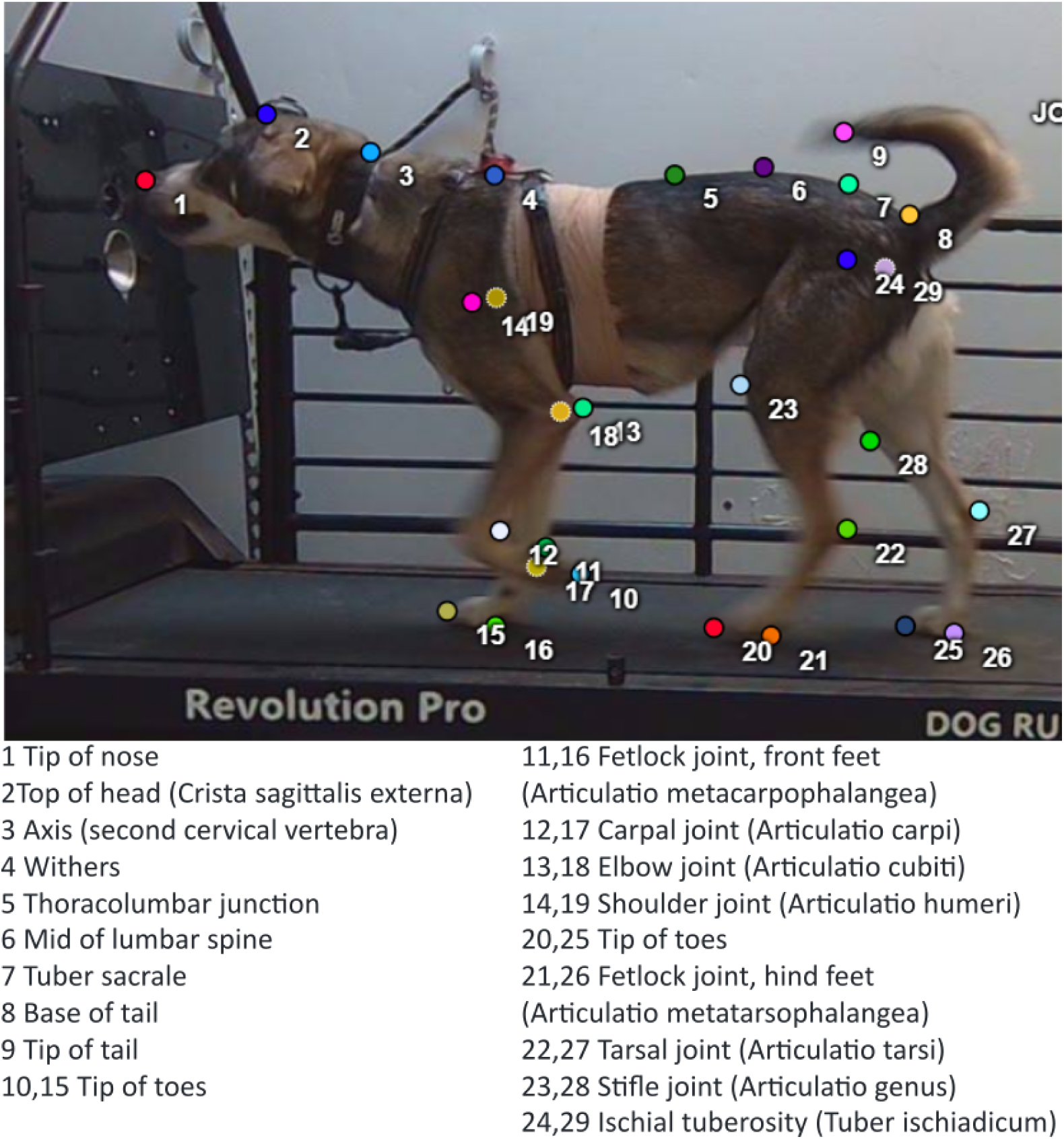
Anatomical landmarks annotated for markerless pose estimation. Twenty-nine landmarks were manually annotated and used for DeepLabCut training, covering cranial reference points, spinal markers, limb joints, and tail positions. Landmark IDs correspond to the anatomical definitions shown alongside the image.

#### Heart rate

Heart rate was sampled at 1 Hz and resampled to yield five equidistant values over the 5-second window. These values were linearly interpolated if necessary to handle occasional missing data (e.g., due to brief loss of electrode contact or movement artifacts). The resulting 5-element vector was used as input to the physiological branch of the multimodal model.

#### Inter-beat interval (IBI)

Inter-beat interval (IBI) data consisted of the time intervals between successive heartbeats, commonly referred to as RR intervals (the time between two consecutive R-peaks on the electrocardiogram). Within each 5-second window, the number of available RR intervals varied depending on heart rate. We therefore computed seven summary statistics: (1) mean RR interval, (2) standard deviation, (3) minimum, (4) maximum, (5) RMSSD (root mean square of successive differences), (6) coefficient of variation, and (7) mean absolute successive difference (MSD). These features capture both average heart rhythm and short-term variability—metrics that are often more sensitive to changes in autonomic state than raw heart rate alone. If fewer than two RR intervals were available, the sample was excluded.

#### Core temperature

Core body temperature was recorded via an ingestible sensor (AniPill™, *manufactured by BodyCap, France*)). For each trial, we selected the most recent temperature measurement at or before odor onset.

All features (pose coordinates, heart rate, IBI statistics, and temperature) were z-scored per dog to remove inter-individual differences in baseline physiology and body size.

### Video-Based Movement Extraction

#### Markerless Pose Estimation

We implemented a two-stage pipeline to extract high-resolution body posture and movement information from video recordings of detection dogs performing odor detection tasks. The first stage used DeepLabCut (DLC) for markerless pose estimation (11), which enables high-precision tracking of user-defined anatomical landmarks without requiring physical markers.

A total of 120 frames were manually annotated using the open-source Computer Vision Annotation Tool (CVAT) (30), selected to span a broad range of postures and movement phases. Annotations included 29 anatomically relevant landmarks, such as cranial reference points (e.g., tip of nose, crista sagittalis), vertebral markers (e.g., thoracolumbar junction), limb joints (e.g., carpal, stifle, tarsal), and the tip of the tail (Figure 1). These landmarks were chosen to provide broad anatomical coverage of the body, with particular emphasis on regions that reflect behaviorally meaningful variation during detection work. Specifically, the selected points capture changes in head posture, gait dynamics, and tail orientation— features known to correlate with odor localization, arousal, and engagement. We further prioritized points that were reliably visible from the lateral camera perspective and consistent across individuals, ensuring robust tracking and cross-dog comparability.

The DLC model was trained on the annotated dataset using standard data augmentation, active learning, and iterative refinement procedures. The final model was then applied to all video frames, generating per-frame pose predictions for each dog using a single model across individuals.

#### Segmentation-Based Refinement of Pose Predictions

To enhance pose estimation accuracy, particularly in regions prone to background clutter or occlusion, we introduced a segmentation-based preprocessing step prior to inference. For each video, we performed an initial DLC prediction on the first frame and selected three stable landmarks along the dog’s torso. These were used to define a region of interest (ROI) centered on the core body.

The ROI was passed to the MediaPipe “Magic Touch” segmentation model (31), which produced a binary mask isolating the dog’s body from the background. This mask was applied to the original frame, and pose estimation was re-run on the segmented image to yield cleaner and more confident landmark predictions.

The segmentation-enhanced procedure was then applied iteratively: for each subsequent frame, the ROI was dynamically updated based on the previous frame’s pose prediction. This allowed for robust tracking without explicit object tracking. The method proved especially effective for improving landmark accuracy on the head and distal limbs, where pose predictions were otherwise affected by background texture or occlusion. All final pose predictions used in downstream modeling were generated from these segmentation-refined frames.

### Model Architecture and Training

#### Spatiotemporal Graph Network (ST-GAT-TCN)

##### Pose-only model

The pose-only version of the model predicts the probability that the dog will alert to the target odor, based solely on the sequence of joint positions extracted from video. Each input sample consists of a sequence of 135 video frames, with 29 anatomical landmarks (joints) per frame. For each joint, we used two-dimensional coordinates (x, y) and a confidence score, resulting in an input tensor of shape (135, 29, 3).

To model spatial structure, we constructed a fixed undirected graph with 29 nodes, where each node corresponds to a joint and edges reflect anatomical connectivity. Edges were defined based on prior knowledge of the canine musculoskeletal system, including linear chains along the spine, symmetric limb segments, and attachments of limbs to the torso. A self-loop was added to every node. The resulting adjacency matrix was used across all time steps.

Spatial processing was performed by a shared two-layer Graph Attention Network (GAT) applied independently to each frame. Each GAT layer used 4 attention heads (concatenated) with 64 output features and ReLU (rectified linear unit) activation. The GAT layers produced a frame-level joint embedding of shape (29, 64), which was then compressed via global average pooling over joints, yielding a single vector per frame.

Temporal modeling was performed by a multi-layer Temporal Convolutional Network (TCN) consisting of 1D convolutional layers with increasing dilation factors [1, 2, 4, 8, 16, 32], a kernel size of 2, and 32 filters. The TCN operated on the time axis and captured behavioral dynamics across the 4.5-second window. The output of the TCN was globally pooled over time and passed through a dense layer (64 units, ReLU activation), followed by a sigmoid output layer predicting the probability of an indication.

The model (Figure 4) was implemented in Python using the Keras framework and the GATConv layer from the Spektral library (32). Code, model weights, and all necessary data are publicly available at https://doi.org/10.5281/zenodo.17197670.

The model was trained using the Adam optimizer with a learning rate of 10*^−^*^3^ (controlling how much model weights are updated in response to errors) and binary cross-entropy loss. Evaluation metrics included binary accuracy, AUC (area under the receiver operating characteristic curve, summarizing classification performance across thresholds), precision, and recall. These metrics were computed across all dogs in the pooled dataset. No explicit dog-level cross-validation was used, as our dataset comprised only four individuals. While a leave-one-dog-out approach would offer stricter generalization testing, it would also reduce the training data per fold to just three dogs, substantially limiting the model’s ability to learn robust spatial–temporal patterns.

Instead, we randomized trial-level assignments across dogs into training, validation, and test sets to evaluate generalization across individuals, while maximizing data availability for learning.

##### Multimodal model with physiological input

To assess whether physiological signals provide complementary information beyond movement alone, we extended the pose-only architecture with three auxiliary input branches for heart rate, inter-beat interval (IBI), and core body temperature. These inputs were processed in parallel to the spatiotemporal pose sequence and fused prior to the classification head.

The physiological input consisted of three components per trial: a 5-element vector of raw heart rate values recorded at 2 Hz (covering approximately 4.5–5 seconds), a 7-element feature vector representing summary statistics of the IBI signal, and a single scalar value representing core body temperature. Each of these inputs was passed through a dedicated small feedforward subnetwork: the heart rate and IBI vectors were each processed by two dense layers (32 and 16 units, ReLU activation), while the temperature input was passed through a single dense layer with 8 units.

The outputs of the three branches were concatenated and projected through an additional dense layer with 16 units (ReLU activation) to form a unified physiological embedding. This embedding was then concatenated with the spatiotemporal pose representation obtained from the ST-GAT-TCN pipeline. The fused feature vector was passed through a final dense classification head identical to the one used in the pose-only model (Figure 4).

This late-fusion design allowed the model to learn from physiological and behavioral features in a complementary fashion, without altering the core architecture of the pose-based pipeline.

#### Training Data and Augmentation

Model training was based on pose sequences collected during treadmill sessions in which dogs were intermittently exposed to target odors. Only trials presenting the target odor were retained, and only if a complete pose sequence was available during the 4.5-second window preceding odor onset. The final training set was constructed from 5-second pose clips (150 frames at 30 Hz), from which we derived both raw and augmented 4.5-second windows.

The base training samples consisted of the final 4.5 seconds (135 frames) of each 5-second clip. To increase training diversity and support generalization, we generated five augmented samples per original trial using a combination of phase shifting and spatial transformations. Specifically, a random 4.5-second window (±15 frames) was extracted from each 5-second clip, and affine transformations were applied to the joint coordinates. These included isotropic and anisotropic scaling, random rotations (±10 degrees), and additive Gaussian noise to the (*x, y*) positions, applied independently per joint and frame.

To mitigate class imbalance, we applied augmentation separately within each class and retained all augmented samples from the minority class. A random subset of augmented samples from the majority class was then added to equalize class sizes. The resulting training set consisted of 1,716 samples, balanced between indication (n = 858) and non-indication (n = 858) trials. Labels were binary: 1 for trials in which the dog correctly indicated the target odor, and 0 for misses.

Augmentation was applied only to the training set. The validation and test sets were held out and included only unaugmented pose sequences.

#### Training Procedure and Evaluation

Each model was trained using the Adam optimizer (*η* = 0.001), binary cross-entropy loss, and early stopping based on the validation AUC (area under the receiver operating characteristic curve). The AUC summarizes the model’s ability to distinguish between positive and negative classes, with 1.0 representing perfect classification and 0.5 indicating chance. It is particularly useful for imbalanced datasets, as it provides a threshold-independent measure of performance that is often more informative than raw accuracy. Training continued for a maximum of 100 epochs, with the best-performing model selected according to the highest validation AUC. To account for variance in network initialization and convergence behavior, each architecture was trained independently three times. In cases where the network failed to escape a local minimum, early stopping typically occurred before epoch 25.

The ST-GAT-TCN network trained on video data only showed some run-to-run variability, with the best model (Trial 1) reaching a validation AUC of 0.882. In contrast, the multimodal model trained on both video and physiological data achieved a substantially higher peak AUC of 0.953 (Trial 3). As shown in Figure 2, the closer match between training and validation curves in this model indicates improved generalization compared to the video-only baseline. Training metrics for all runs are summarized in Table 2.

**Fig 2.**
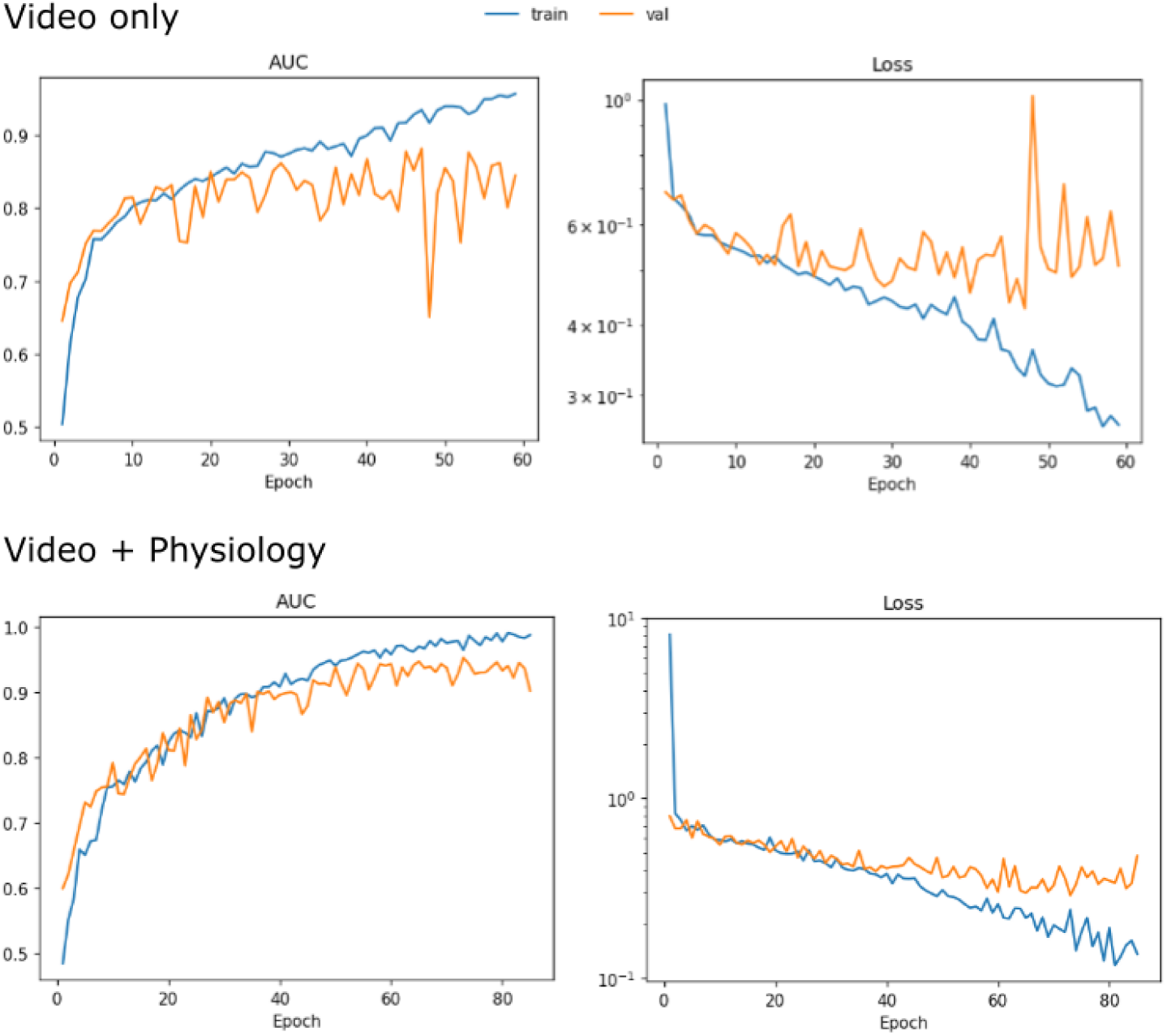
Training dynamics of the video-only and multimodal models. Top row: performance of the ST-GAT-TCN model trained solely on video-based joint trajectories. Bottom row: performance of the multimodal model with additional physiological inputs (heart rate, inter-beat interval, and temperature). Left column shows AUC; right column shows binary cross-entropy loss. Curves are shown for the best-performing run of each model. Blue lines (“train”) show training metrics. Orange lines (“val”) show validation metrics. Note the stronger alignment between training and validation curves in the multimodal model, suggesting improved generalization. Full metric trajectories (including accuracy, precision, and recall) are provided in Supplementary Figures S1–S2.

**Table 2.** Validation metrics at best epoch for each training run. Bold indicates the best value across all runs per column (excluding epoch). Abbreviations: AUC = area under the ROC curve, Acc = accuracy, Loss = binary cross-entropy loss, Prec = precision, Rec = recall.

| Trial | Epoch | AUC | Acc | Loss | Prec | Rec |
| --- | --- | --- | --- | --- | --- | --- |
| **Video only** |  |  |  |  |  |  |
| 1 | 47 | 0.882 | 0.798 | 0.427 | 0.824 | 0.763 |
| 2 | 16 | 0.831 | 0.743 | 0.511 | 0.710 | 0.786 |
| 3 | 17 | 0.847 | 0.763 | 0.493 | 0.718 | **0.840** |
| **Video + physiology** |  |  |  |  |  |  |
| 1 | 20 | 0.872 | 0.769 | 0.466 | 0.845 | 0.665 |
| 2 | 19 | 0.864 | 0.790 | 0.467 | 0.788 | 0.795 |
| 3 | 73 | **0.953** | **0.870** | **0.290** | **0.899** | 0.844 |

## Results

### Segmentation Enhances Signal-to-Noise Ratio of Markerless Pose Estimation

The predictive framework we developed relies critically on accurate and consistent pose estimation from video. Any errors in locating joints could mask the subtle movement patterns we aim to detect and weaken subsequent model performance. However, achieving this accuracy is challenging in our olfacto-treadmill recordings, where fixed background elements—such as the treadmill frame and the dark panel housing of the olfactometer—introduce strong visual edges and textures that can confuse the model. These structures can resemble parts of the dog in shape or motion, leading to reduced confidence and precision in joint localization. To address this, we evaluated whether a segmentation-based preprocessing step, designed to isolate the dog from its surroundings, could improve the quality of pose predictions and thereby strengthen the signal available for downstream behavioral analysis.

We applied a lightweight enhancement pipeline that combined pose prediction with background removal (see Materials and Methods and Supplementary Video S1). In each frame, we first defined a region of interest (ROI) around the dog’s torso based on the previous frame’s joint positions. This ROI was passed to a segmentation model, which isolated the dog from the background before re-running pose estimation on the segmented image.

To quantify the effect of segmentation, we compared DeepLabCut likelihood values between the standard (“full”) pipeline and the segmentation-enhanced (“segmented”) version. Averaging over all keypoints and test frames, pre-segmentation yielded significantly higher likelihoods (U = 48,421, *p <* 0.00001; Figure 3, Panel A). For keypoints with likelihood 0.6, segmentation also reduced the mean pixel-wise error from 20.53 to 7.11 pixels, indicating improvements in both confidence and localization accuracy.

**Fig 3.**
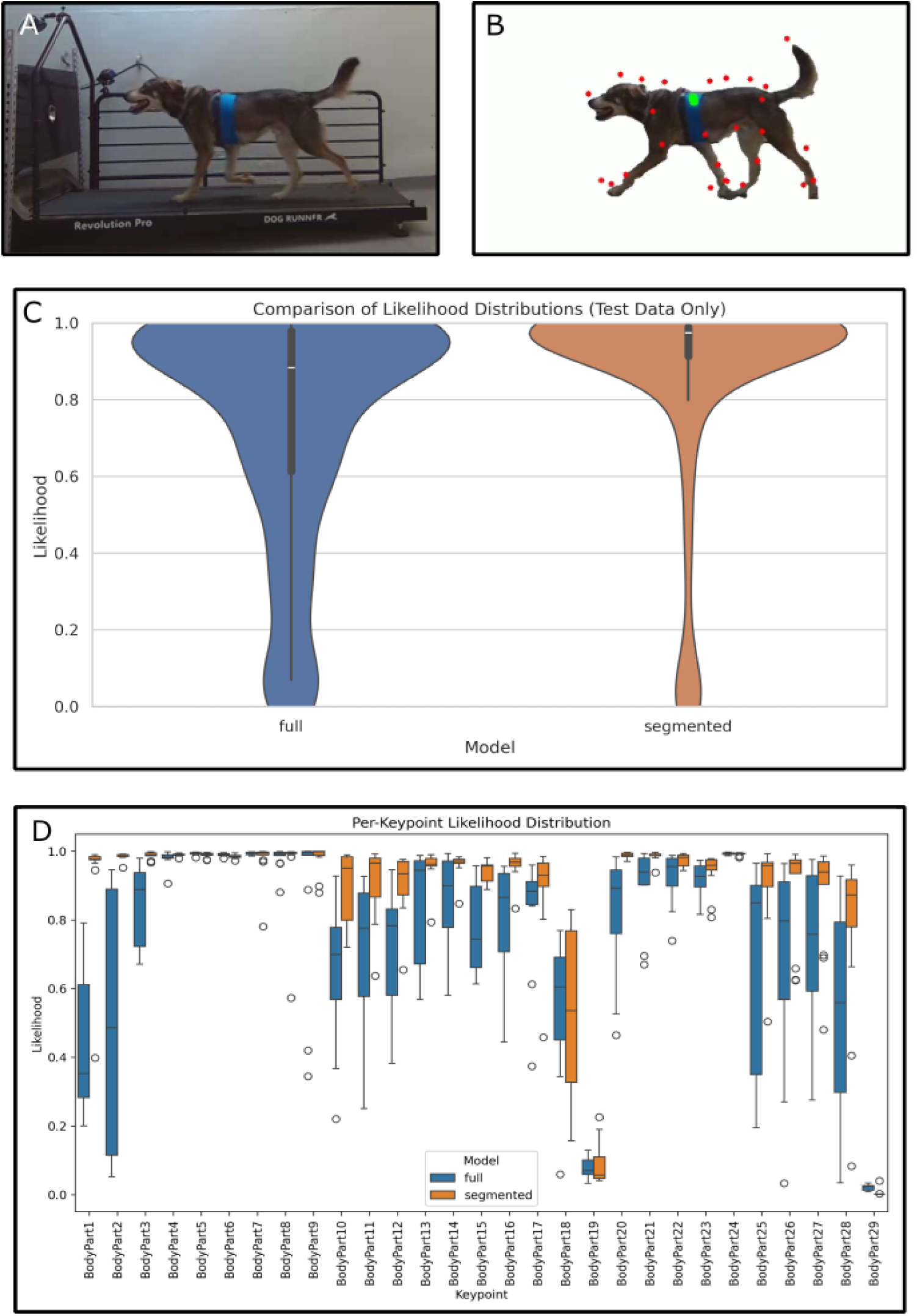
Segmentation improves pose prediction quality by enhancing signal-to-noise ratio. (**A**) Sample frame from the olfacto-treadmill recording setup. Dogs performed repeated trotting trials while being exposed to controlled odor pulses. (**B**) Output of the segmentation-enhanced pose prediction pipeline. Joint markers are overlaid on the segmented frame. Background elements such as the treadmill structure are effectively removed, reducing visual clutter. (**C**) Violin plots comparing DeepLabCut likelihood distributions between the full (unsegmented) and segmented pipelines on the test set. Segmentation significantly increased pose confidence (Wilcoxon U test, *p <* 0.00001). (**D**) Per-joint boxplots showing likelihood distributions for each keypoint. Gains were most prominent in the head, cranial spine, and distal joints. Full statistical results are available in Supplementary Table S1.

Analysis of per-joint likelihood distributions (Figure 3, Panel B) showed that these benefits were not uniform across the body. Significant improvements (*p* ¡ 0.05, Wilcoxon U test) were most pronounced in the head region (BodyParts 1–4), cranial spine (BodyParts 2–3), and distal limb joints (e.g., 10–12, 20–23) (Supplementary Table 1). Together, these results demonstrate that pre-segmenting the image enhances pose prediction quality in terms of both confidence and spatial precision. The particularly marked gains in cranial landmarks—highly relevant in olfactory tasks—suggest that segmentation may be especially valuable for studies focusing on head posture and orientation in odor-based work.

### Graph-Temporal Network to Capture Structure and Dynamics

With the segmentation step improving the precision and confidence of pose estimates, we next turned to the question of how best to interpret these joint trajectories to predict performance outcomes. Reliable localization alone is insufficient; the model must also capture the structural relationships between joints within each frame and the temporal patterns of movement across frames. To address this, we developed a two-stage architecture that combines graph-based reasoning with temporal pattern recognition (Figure 4).

**Fig 4.**
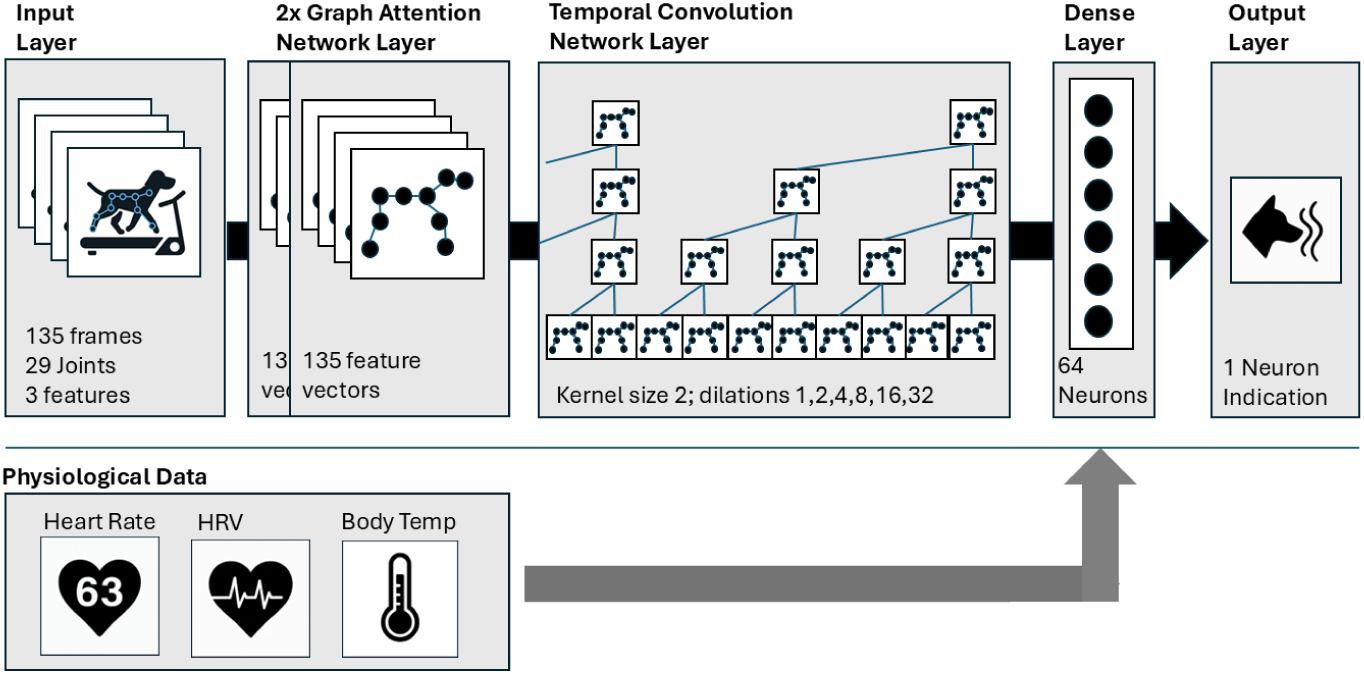
Architecture Spatio-Temporal Network Models. The network processes 135-frame sequences of 29 joints (with x, y, and confidence as features). Pose graphs from each frame are first passed through two stacked Graph Attention Network (GAT) layers, enabling information exchange up to the second-order neighborhood of each joint. These spatial embeddings are then processed by a multi-layer Temporal Convolutional Network (TCN) with increasing dilation factors to capture movement dynamics over time. A final dense layer maps the sequence representation to a binary output predicting whether the dog will indicate a target odor. In an extended version of the model, physiological data (heart rate, heart rate variability, and core temperature) are processed through separate dense layers and fused with the pose-based representation before the final dense layer.

In each frame, the predicted body keypoints form a structured representation of posture. Rather than treating these coordinates as an unstructured list, we embedded them in a graph in which nodes represent joints and edges connect anatomically adjacent joints. This preserves the body’s structural topology and allows the network to model joint–joint dependencies explicitly. To process this graph, we employed a Graph Attention Network (GAT) (33). Each GAT layer enables a joint to “attend to” its immediate anatomical neighbors, dynamically weighting their contribution to the current representation. Stacking two GAT layers allows each joint’s representation to be influenced not only by its direct neighbors but also by second-order (2-hop) connections, thereby capturing both localized configurations and broader relational patterns across the skeleton. Technically, the graph-based encoder processes each frame independently, producing a single vector embedding of the dog’s posture via two stacked GATConv layers (each with four attention heads and 64 hidden units), followed by a global average pooling step across joints.

Since our goal was to analyze movement rather than static posture, it was essential to model the temporal evolution of these embeddings. To achieve this, we passed the per-frame graph embeddings through a Temporal Convolutional Network (TCN), a specialized architecture for sequence modeling that uses layers of dilated one-dimensional convolutions (34). This design allows the network to detect movement motifs—such as head dips or changes in stride regularity—across multiple timescales. We implemented six dilation layers with receptive fields increasing from 1 to 32 frames, enabling the model to integrate both short- and long-range temporal dependencies.

The complete architecture—comprising the per-frame graph encoder and the TCN sequence processor—was trained end-to-end. The temporally processed features were then passed through a final dense layer to predict whether the dog would indicate a target odor after the given movement segment. In total, the model contained approximately 50,000 trainable parameters.

### Model Learns to Predict Missed Indications from Video Alone

Having established the architecture, the next step was to evaluate whether it could detect the subtle movement signatures that precede a missed indication. To ensure that any predictive information came from the dog’s ongoing behavior rather than a direct reaction to odor, we trained the model exclusively on pose sequences from the 4.5 seconds immediately before odor presentation. This design tests the model’s ability to anticipate performance outcomes based solely on pre-stimulus movement patterns.

The model was trained on a balanced, augmented dataset derived from treadmill trials (see *Materials and Methods*). Each trial provided a sequence of 135 frames, with 29 joints per frame and three features per joint (x, y, likelihood). The architecture processed each sequence end-to-end, outputting a binary prediction: indication (1) or non-indication (0).

Training used an 80/20 split for training and validation. We optimized with Adam (learning rate = 0.001) and binary cross-entropy loss, tracking binary accuracy, area under the ROC (receiver operating characteristic) curve (AUC), precision, and recall as performance metrics. Early stopping was applied with validation AUC as the primary criterion, halting if no improvement greater than 0.001 was observed over 12 consecutive epochs. To mitigate variability in convergence—likely due to sensitivity to initialization and data ordering—we trained three independent runs with different random seeds, selecting the model with the highest validation AUC for final evaluation.

The selected model converged after 47 epochs, achieving a validation AUC of 0.8820, accuracy of 0.7983, precision of 0.8238, and recall of 0.7626. These results indicate that the network successfully captured meaningful spatial–temporal patterns predictive of trial outcomes.

On the held-out test set of 216 trials (144 non-indications, 72 indications), the model achieved an overall accuracy of 73.6% and a AUC of 0.81, demonstrating good discriminative capacity. Performance was particularly strong for the non-indication class—often the more operationally relevant case—reaching a precision of 0.82 and a recall of 0.77. This means the model correctly identified most trials in which the dog did not respond, while maintaining a low false-alarm rate. For the indication class, performance was lower (precision = 0.59, recall = 0.67), indicating that some true indications were missed or misclassified.

Overall, these findings confirm that reliable predictive information is embedded in the dog’s movement patterns and posture before encountering the target odor. The model’s strong performance on non-indications suggests particular promise for early detection of performance decline, where timely intervention could prevent a critical miss during operational deployment.

### Physiological Signals Enhance Prediction Based on Video

The video-only model demonstrated that a 4.5-second window of pre-odor movement contains sufficient information to predict whether a dog will indicate a target odor. Building on this finding, we asked whether predictive performance could be further enhanced by incorporating additional modalities. Physiological signals were a logical choice, as they provide direct measures of the dog’s internal state that are not captured in posture or gait. We therefore extended the model to include three well-established indicators in canine performance research: heart rate (HR), heart rate variability (HRV, quantified via inter-beat interval), and core body temperature (Temp).

To implement this extension, we modified the original architecture to incorporate the three physiological inputs recorded during the same pre-odor window. Each modality was processed through a dedicated dense branch, and the resulting embeddings were concatenated with the pose-based representation prior to the final classification layer (see *Materials and Methods* for architectural specifics). The input for each sample thus comprised (i) a 135-frame pose sequence across 29 joints with three features per joint (x, y, likelihood), (ii) a 9-element vector summarizing HR in the window, (iii) a 7-element vector summarizing IBI/HRV (e.g., mean RR and time-domain variability measures), and (iv) a single scalar for Temp. This multimodal design enabled the network to jointly model external movement patterns and internal physiological state.

As with the video-only model we trained three independent instances of the multimodal model (different random seeds) using the same procedure as the video-only baseline: an 80/20 split of the augmented, class-balanced dataset; Adam optimizer (learning rate = 0.001); binary cross-entropy loss; and monitoring of accuracy, ROC AUC, precision, and recall on both training and validation sets. Early stopping was applied based on validation AUC (patience = 12, AUC 0.001), with restoration of the best weights. Among the three runs, the best-performing multimodal model achieved a validation AUC of 0.9534, exceeding the best video-only model (0.8820). Validation accuracy (0.8696), precision (0.8985), and recall (0.8442) were correspondingly high, with no evidence of overfitting (Table 2).

On the held-out test set (216 trials: 144 non-indications, 72 indications), the multimodal model achieved an overall accuracy of 76.85% and a ROC AUC of 0.80. For the non-indication class (class 0), precision was 0.81 and recall was 0.85. The corresponding F1-score—defined as the harmonic mean of precision and recall—was 0.83. This indicates that most missed indications were correctly identified with few false alarms—an operationally desirable balance. Performance on the indication class (class 1) was lower (precision = 0.67, recall = 0.61), consistent with a model tuned more conservatively to avoid missed non-indications. Relative to the video-only baseline, the inclusion of physiological data yielded a recall gain for non-indications of 0.85 vs. 0.77, with precision essentially unchanged, resulting in a higher F1 for class 0 (Figure 5).

**Fig 5.**
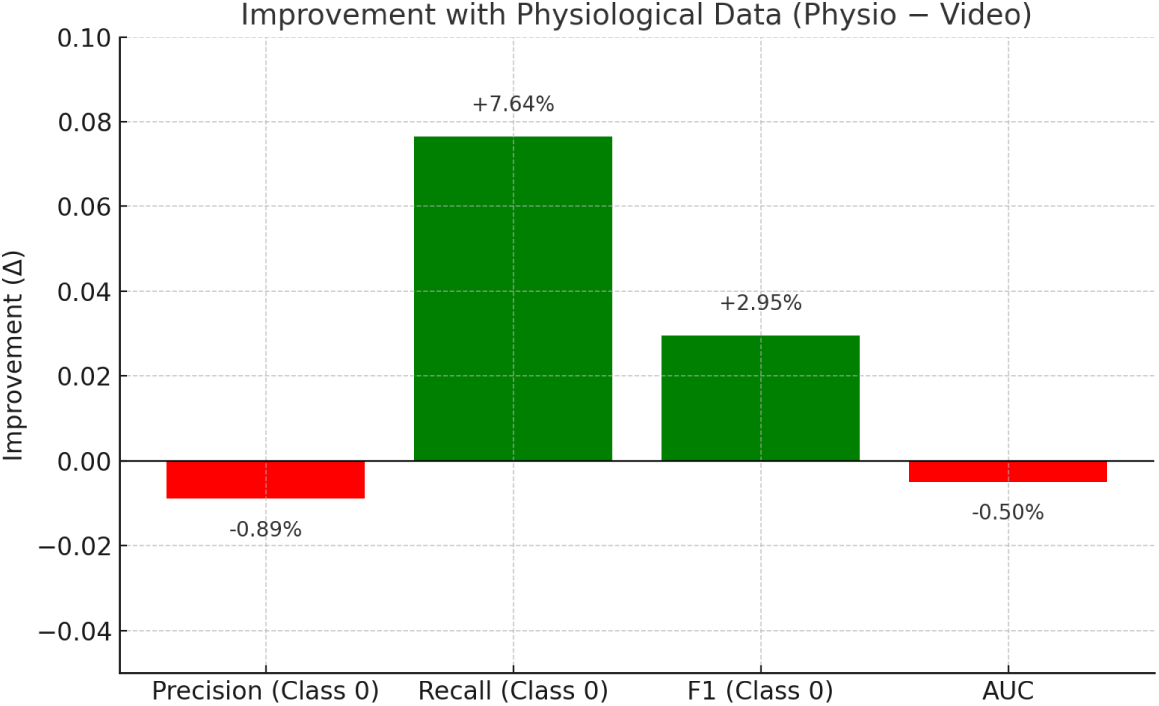
Improvement in model performance for predicting missed indications when physiological data (heart rate, heart rate variability, and core temperature) were included alongside video-derived pose features. Bars show the change in key evaluation metrics for the no-indication class (class 0) on the held-out test set. While precision remained nearly unchanged, the model with physiological data achieved higher recall and F1-score, correctly identifying more missed indications without sacrificing precision. This suggests that physiological inputs provide additional predictive value beyond movement data alone.

The integration of HR, HRV (IBI), and Temp provides complementary information beyond movement alone, improving sensitivity to trials that culminate in a missed indication without inflating false positives. This supports the use of multimodal sensing to monitor performance readiness in detection dogs and motivates downstream analyses to determine which physiological inputs contribute most strongly to the observed gains.

When compared to the model trained on video data alone, the model incorporating physiological information showed consistent advantages in detecting non-indication trials (Figure 5). While both models achieved similar precision for class 0 (0.82 for video-only vs. 0.81 for video+physio), the inclusion of physiological data led to a marked increase in recall (0.85 vs. 0.77), resulting in a higher F1-score (0.83 vs. 0.80). This indicates that the multimodal model was more sensitive to relevant behavioral and physiological cues associated with missed indications, without increasing the rate of false positives.

In operational terms, this improvement means that the combined model is more reliable in correctly identifying trials where the dog does not respond to the target odor. This is particularly important for early detection of performance decline or fatigue, where missed indications may signal that the dog is not fully engaged or in an optimal physiological state. The integration of heart rate dynamics and core body temperature appears to provide valuable information about the dog’s internal state, complementing the observable movement patterns captured in the video data.

### Explaining Model Decisions: Which Inputs Matter Most?

The results above show that combining movement and physiological signals enables our model to reliably anticipate missed indications. While these performance metrics demonstrate that both modalities carry predictive value, they do not explain which parts of the input have the greatest influence on the model’s predictions. For practitioners—particularly dog handlers and veterinarians—this is a critical question: if a predictive system is to be trusted and acted upon, its attention should align with cues that experienced handlers themselves consider meaningful. Identifying these influential features can also offer novel insights into behavioral and physiological markers of fatigue or disengagement, potentially informing future training or monitoring strategies.

To explore this, we examined the model’s internal focus in two complementary ways.

First, for the video data, we asked which body parts most strongly influenced its decisions, effectively mapping its attention across the dog’s posture and movement. Second, for the physiological data, we tested the impact of removing each signal in turn and observed how much predictive performance declined. Together, these analyses reveal not just that the model works, but how it works—linking its decisions to tangible, real-world cues. This provides a bridge between the statistical performance measures of the previous sections and the practical knowledge base of experienced handlers, and may guide future development of wearable sensors and behavioral assessment tools for operational use.

#### Which joints matter: Saliency analysis of pose-based predictions

To identify which anatomical landmarks most strongly influenced the model’s predictions, we analyzed the video-only model using a gradient-based method that estimates the impact of small changes in each joint’s predicted position or detection confidence on the model output (35). For every trial in the test set, we computed these values for all joints, averaged them across frames and trials, and rescaled them by a factor of 100 for readability. Higher values indicate joints whose variation had a greater effect on the predicted outcome.

The resulting ranking (Figure 6) shows a clear pattern. Cranial landmarks dominate the profile: the top of the head (0.199), the second cervical vertebra (C2, 0.09), and the tip of the nose (0.059) exerted the strongest influence, reflecting subtle changes in head posture—such as nose dips or freezes—often associated with alert behavior. The tip of the tail (0.085) also ranked highly, suggesting that tail movement provides information relevant to the dog’s behavioral state. An unexpected outlier was the right ischial tuberosity (0.162), which is not directly visible in the video and receives zero likelihood in most frames. This is likely a gradient artifact caused by numerical instability in the saliency calculation for an occluded joint, and should therefore be interpreted with caution. Other joints, including distal limbs and thoracolumbar landmarks, had considerably lower values (*<* 0.033), indicating a smaller direct influence on the model’s decisions.

**Fig 6.**
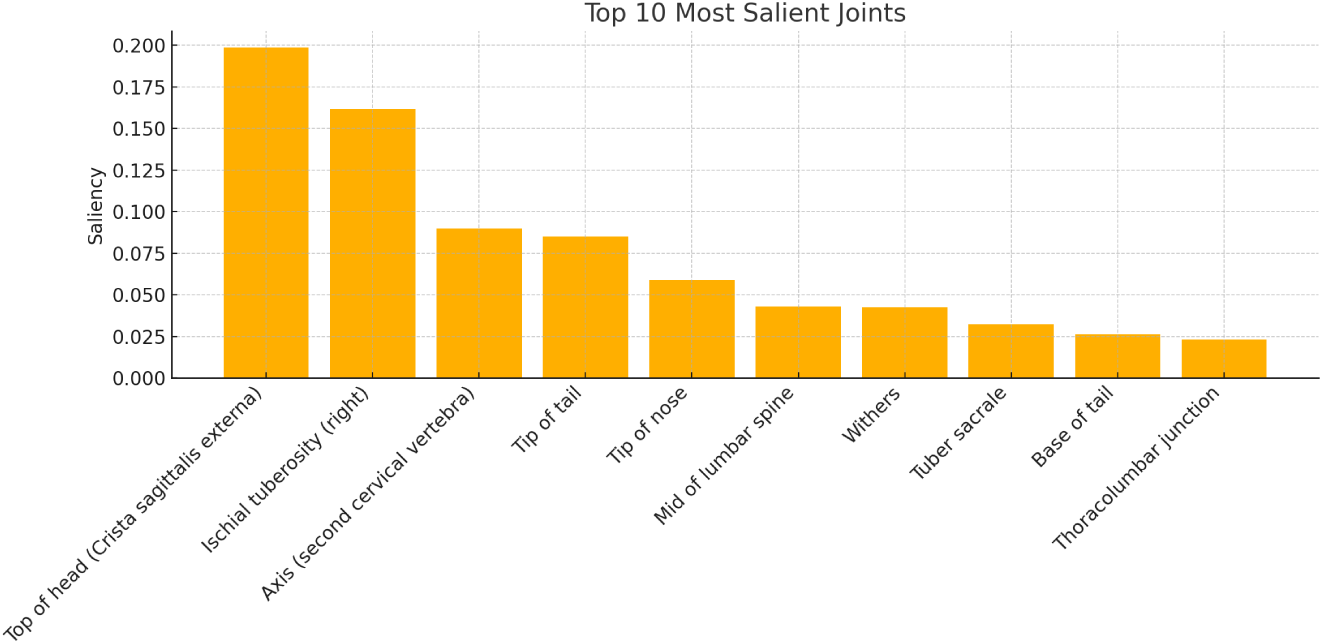
Top 10 most salient joints based on gradient analysis of the video-only model. Saliency values were averaged over all frames and all test trials, and rescaled by a factor of 100 for readability. Cranial landmarks (top of head, axis, tip of nose) and the tip of the tail show the strongest influence on the model’s prediction. The high saliency of the right ischial tuberosity likely reflects a gradient artifact, as this joint was consistently occluded in the video data (see Supplementary Video S2 for frame based analysis).

Because our architecture includes two stacked Graph Attention Network layers, each joint’s representation incorporates information from both its direct anatomical neighbors and from joints two steps away in the skeletal graph. High saliency at a particular joint therefore does not imply that the model uses that joint in isolation; rather, it likely reflects the importance of the broader neighborhood in which it is embedded.

In summary, the model’s predictions depend most heavily on a small set of cranial and caudal landmarks. This pattern points to head posture and tail behavior as key contributors—features that experienced handlers often monitor during odor localization and alerting.

#### Which physiological signals matter: Sensor ablation analysis

Given the performance gains from including physiological inputs (Section 3.4), we next examined which individual signals contributed most to the model’s predictions. Identifying these contributions is important not only for understanding the model’s internal logic, but also for clarifying which physiological processes may underlie performance decline and which sensors would be most valuable for field deployment.

We conducted a sensor ablation study (36) in which each of the three physiological branches—heart rate, inter-beat interval, and core body temperature—was removed in turn by replacing its input values with zeros, while keeping all other data unchanged. This procedure mimics a scenario where a sensor is unavailable due to malfunction or omission. Predictions were then recomputed on the held-out test set and compared against the full-input condition AUC, accuracy, and log-loss.

The ablation results showed clear differences in importance between signals (Table 3). Removing the inter-beat interval branch caused the largest degradation in performance. AUC dropped by 0.16 (95% CI: 0.08–0.23), accuracy fell by 6 percentage points (*p* = 0.13), and log-loss increased by 0.19 (*p* = 0.035). While not all effects reached statistical significance, the consistent performance drop across metrics identifies inter-beat interval as the most critical physiological signal for predicting missed indications. Eliminating core temperature produced a moderate but significant drop in performance (AUC = 0.06, 95% CI: 0.03–0.10; Accuracy = –4.1%; Log-loss = 0.20, *p <* 0.001), indicating that the model also exploits subtle thermal deviations, albeit less strongly than heart rate variability.

**Table 3.** Physiological signal ablation effects on model performance. Arrows indicate direction and relative magnitude of change vs. full model (↓↓↓ = large drop; ↓ = small drop; ≈ = no change; ↑ = increase). *^†^* = significant (*p <* 0.05); n.s. = not significant.

| Ablation | Rank (AUC) | Acc | Log-loss | Interpretation |
| --- | --- | --- | --- | --- |
| No IBI | $\downarrow\downarrow\downarrow^\dagger$ | ↓ n.s. | $\uparrow^\dagger$ | Most critical physiological input |
| No Temp | $\downarrow^\dagger$ | ↓ n.s. | $\uparrow^\dagger$ | Moderately informative |
| No HR | ≈ n.s. | ≈ n.s. | $\uparrow\uparrow^\dagger$ | Mainly affects calibration |

In contrast, suppressing heart rate had no adverse effect on AUC or accuracy; the slight AUC increase was not significant, and classification performance remained stable. However, log-loss increased markedly (Log-loss = 0.49, *p* ¡ 0.001), suggesting that heart rate contributes primarily to probability calibration rather than to the decision boundary itself.

Overall, these findings indicate that short-term heart rhythm variability provides the strongest physiological cue for early detection of performance decline, followed by core temperature. Heart rate alone, although traditionally monitored in working dogs, appears to add limited discriminative value in this context.

### From Model Predictions to Canine State Estimation

#### Bayesian integration of model confidence: estimating latent miss-rate

The preceding analyses demonstrate that our model can predict, with reasonable accuracy, whether a detection dog will miss an indication based on short segments of pre-stimulus video and physiological data. Importantly, these predictions are not arbitrary: saliency and ablation analyses confirm that they are driven by meaningful behavioral and physiological features. However, in operational contexts, decisions are rarely made on the basis of isolated predictions. Rather than responding to individual moments, experienced handlers form a continuous, evolving assessment of the dog’s engagement and reliability.

This distinction has important implications for the application of model outputs.

The critical question in practice is not merely whether a given trial will result in a miss, but whether the dog is currently operating in a state where the likelihood of missing a target odor is elevated—e.g., above a context-specific threshold such as 10%. This reframing shifts the focus from binary classification to the estimation of an underlying, latent performance state.

This conceptual shift—from predicting single trial outcomes to estimating a latent performance state over time— naturally lends itself to a Bayesian framework. Such an approach allows probabilistic evidence to be integrated across trials, incorporates prior knowledge about the dog or task, and produces interpretable estimates that support threshold-based decision-making. Critically, it aligns the statistical logic of model aggregation with the cognitive strategies employed by experienced handlers, who assess behavioral trajectories rather than reacting to isolated events.

Implementing such a Bayesian framework, however, typically requires well-calibrated model outputs—i.e., confidence scores that correspond to the true probability of a missed indication. To meet this requirement, we applied post-hoc calibration using temperature scaling (37). This procedure improved several standard calibration metrics. The Brier score (which measures the mean squared difference between predicted probabilities and actual outcomes) decreased from 0.183 to 0.169. The LogLoss (a measure of confidence-weighted error) dropped from 0.699 to 0.515 and the Expected Calibration Error (ECE)—which quantifies the mismatch between predicted probabilities and observed frequencies—was reduced from 0.134 to 0.065.

Despite the improvements in calibration metrics, the process introduced new challenges. Temperature scaling reduced the dynamic range of the model’s predictions: lower scores were inflated (e.g., 0.063 became 0.277), while higher scores were pulled down (e.g., 0.954 became 0.747). As a result, the outputs became less interpretable for end-users—especially in operational contexts where extreme values may carry more weight. This compression also reduced the model’s sensitivity to subtle behavioral shifts, which are often the earliest signs of fatigue or disengagement. Finally, because of the limited size and imbalanced label distribution in our test set, the calibration itself may be unstable, raising caution about its generalizability.

Given these constraints, we opted not to rely on formally calibrated probabilities.

Instead, we used the model’s raw output scores *c_t_* ∈ [0, 1] as internally consistent indicators of certainty. While suboptimal from a probabilistic standpoint, these raw scores preserved ordinal information and relative signal strength—properties sufficient for tracking trends over time. We therefore used them as soft evidence within the Bayesian framework, enabling a continuous estimate of the dog’s latent miss-rate that evolves with each new trial.

To operationalize this approach, we defined a probabilistic model in which the dog’s current miss-rate *θ* ∈ [0, 1] is treated as a latent variable. The model’s raw output scores *c_t_* ∈ [0, 1] are interpreted as fractional evidence for a miss and are integrated over time to update our belief about *θ*. Specifically, we model:

- A prior over *θ* as a Beta distribution:

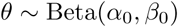

where the mean 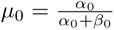 encodes prior expectations about the dog or task, and the total count *κ* = *α*_0_ + *β*_0_ represents prior certainty.

- A sequence of observations {*c*_1_*, c*_2_*, . . ., c_T_* }, each interpreted as soft confidence that a miss will occur on trial *t*.
- The posterior distribution after *T* trials:

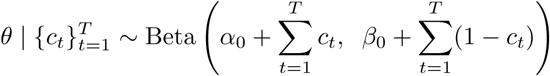

This posterior enables us to compute the probability that the dog’s miss-rate currently exceeds a decision-relevant threshold, for example:

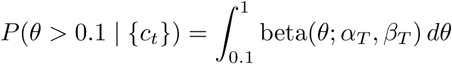

with *α_T_*= *α*_0_ + Σ*c_t_*, and *β_T_* = *β*_0_ + Σ(1 − *c_t_*).

This formulation enables a principled estimate of how likely it is that the dog is currently in a high-risk state— precisely the kind of information that supports real-world deployment decisions.

#### From theory to practice: estimating state in real trial sequences

To illustrate the practical applicability of the Bayesian integration framework, we analyzed a complete run from a single dog performing a standard search task. The run comprised 100 trials, of which 50 included a target odor presentation. For each of these 50 target-present trials, the posterior distribution over the miss-rate *θ* was computed using only the model outputs from a fixed-length observation window preceding odor presentation. From this posterior, we derived the probability that the dog’s miss-rate exceeded two operationally relevant thresholds: 50% and 60%, denoted *P* (*θ >* 0.5) and *P* (*θ >* 0.6), respectively.

These probabilities do not represent the observed miss-rate directly, but rather the posterior estimate that the dog was operating in a state where more than half (or more than 60%) of targets would be missed if the current internal state persisted.

To compare these estimates with actual performance, we computed a ground-truth sliding-window miss-rate across sequences of 10 consecutive target-present trials. As shown in Figure 7, the posterior probabilities (green and blue lines) closely tracked the observed miss-rate (orange), increasing in parallel as the dog began to accumulate missed trials.

**Fig 7.**
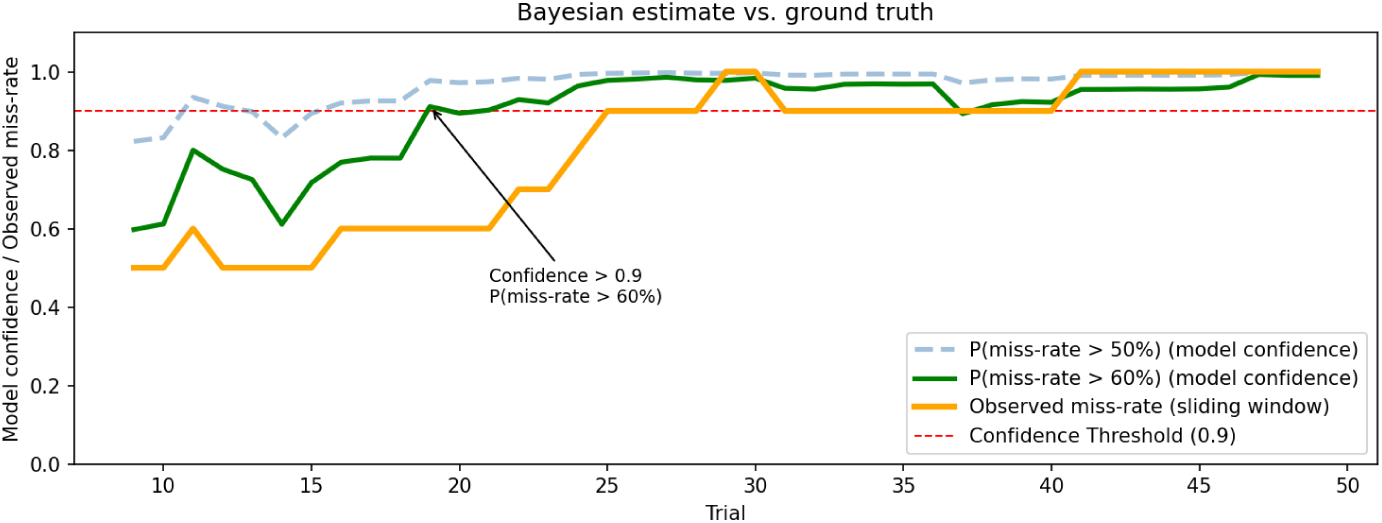
Bayesian aggregation of model confidence over a full target run. For each of 50 target-present trials, we compute the model’s confidence that the dog’s miss-rate exceeds 50% (blue, dashed) or 60% (green), based on pre-odor video and physiological data. These confidence values are derived from the posterior distribution over the latent miss-rate *θ* . The orange line shows the actual fraction of missed indications computed in a 10-trial sliding window. The red dashed line marks a fixed confidence threshold (0.9) for visual reference. The model predictions align closely with the dog’s observed performance, despite having no access to trial outcomes.

Importantly, the model had access only to video and physiological data recorded *prior* to odor presentation and thus no direct information about trial outcomes. The close alignment between posterior estimates and behavioral changes indicates that the aggregated model outputs capture pre-indication patterns predictive of declining performance.

The example run was drawn from the training set and selected for its completeness and continuous temporal context. While this makes it well suited for illustrating the method, validation on fully held-out sequences will be necessary to assess generalization in operational settings.

A key strength of the Bayesian framework is its ability to incorporate prior knowledge about the expected difficulty of the task or the individual dog’s baseline performance. In the example above, we used a neutral prior with *α*_0_ = *β*_0_ = 1, corresponding to an expected miss-rate of 50% with low certainty—reflecting the experimental design, in which odor concentrations were titrated to yield approximately 50% misses. Different priors allow rapid adaptation to other contexts: for a more difficult odor or lower concentration, *α*_0_ = 0.5*, β*_0_ = 1.5 shifts the prior expectation toward 75% misses; for an easier task or highly trained dog, *α*_0_ = 1.5*, β*_0_ = 0.5 centers the prior at 25%. The strength of the prior, given by *α*_0_ + *β*_0_, controls how strongly these expectations influence the posterior; for example, *α*_0_ = *β*_0_ = 5 encodes a strong belief in a balanced 50% miss-rate, making the posterior less sensitive to short-term fluctuations in model output. This property is particularly useful when working with dogs known to be consistent across sessions or when early predictions are noisy.

Figure 8 illustrates how both the prior mean and strength affect the posterior probability that the current miss-rate exceeds 60%, given the same sequence of model outputs. This flexibility enables the framework to be adapted to dog-specific and task-specific operational requirements.

**Fig 8.**
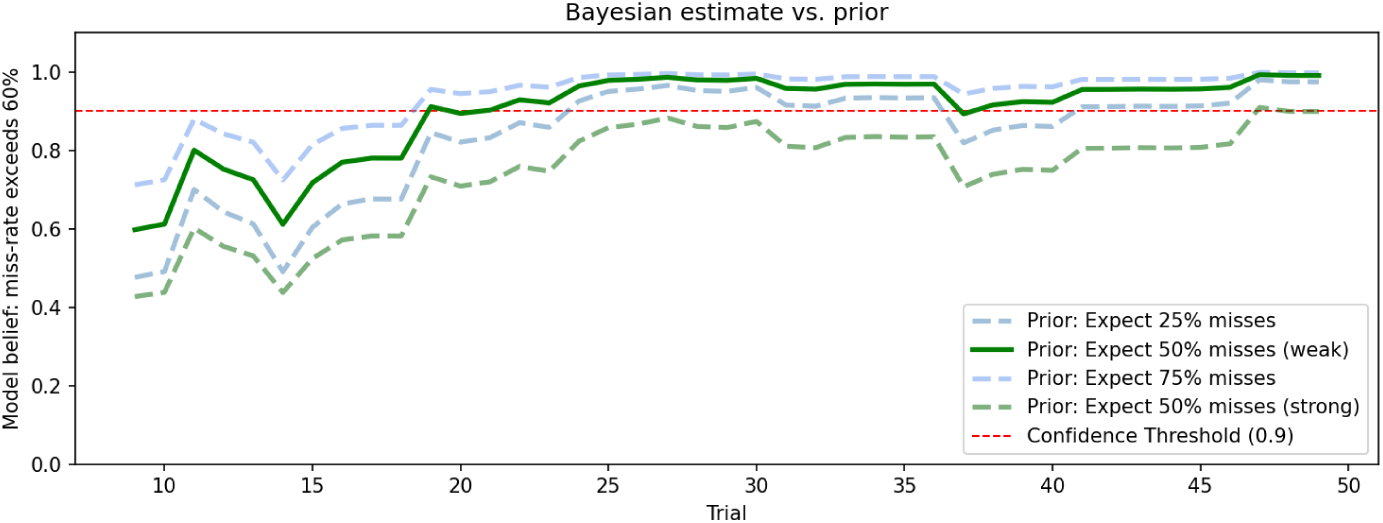
Influence of prior assumptions on Bayesian state estimation. The figure shows the estimated probability that the dog’s current miss-rate exceeds 60%, computed from the same model outputs but under different prior assumptions. The green line corresponds to a weakly informative neutral prior (expected miss-rate 50%, *α*_0_ = *β*_0_ = 1) . Blue and light-green dashed lines show priors with shifted expectations (25%, 75%), while the dark green dashed line represents a strong neutral prior (*α*_0_ = *β*_0_ = 5) . The red dashed line marks a fixed threshold (0.9) for reference. Stronger priors result in more conservative updates, while different prior expectations allow task-specific adaptation.

## Discussion

### Modeling the Missed Alert: A Deep Learning Approach to Canine Performance Prediction

Detection dog performance is mission-critical in many operational settings, from routine inspections to high-stakes deployments such as explosives detection—where a single missed alert can carry severe consequences. Handlers are therefore trained to closely observe their dogs for any signs of declining performance. Yet especially in operational environments, recognizing these early cues can be challenging: distractions, limited visibility, and competing demands may all interfere with continuous monitoring. At the same time, experimental studies—including our own—have shown that detection performance can decline even in the absence of overt behavioral changes. This creates a critical gap: early internal signs of fatigue or disengagement may go unnoticed until a miss occurs.

In this study, we asked whether such early warning signals could be identified using short pre-stimulus segments of movement and physiology recorded under controlled treadmill conditions. By applying a deep learning model to these data and embedding its predictions within a Bayesian decision-support framework, we were able to estimate the evolving likelihood that a dog is operating in a high-risk state for missed indications (Figure 9). The goal was not merely to flag individual trial outcomes, but to track readiness over time in a way that reflects how experienced handlers assess canine performance in the field.

**Fig 9.**
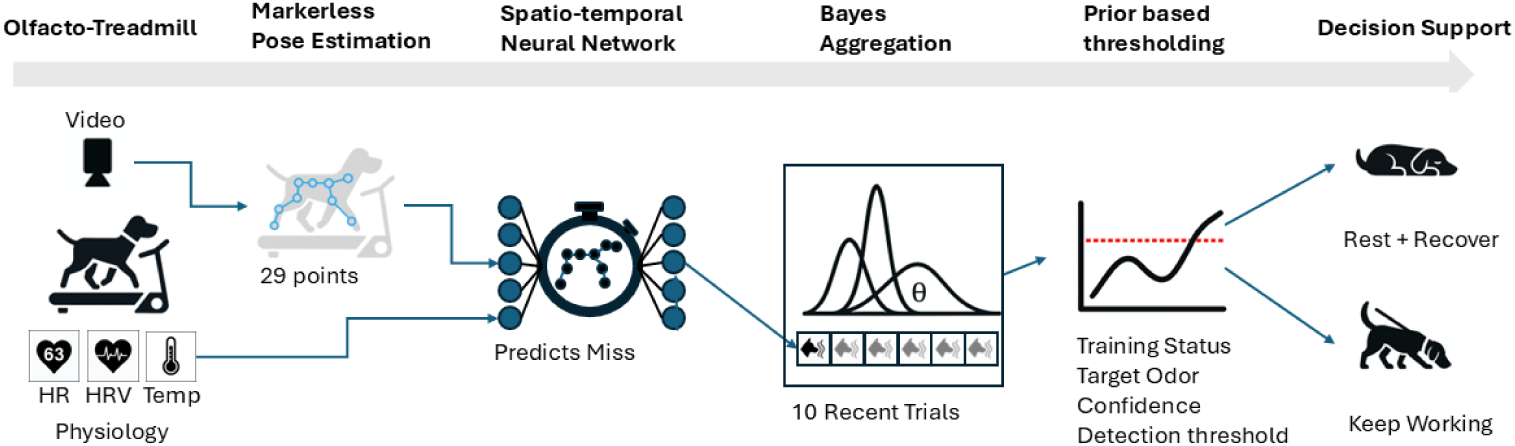
End-to-end system for predicting detection dog performance declines. Video recordings and physiological signals (heart rate, heart rate variability, and core body temperature) are collected during olfacto-treadmill sessions. A markerless pose estimation system extracts movement features, which are combined with physiological input and processed by a spatio-temporal neural network trained to predict missed indications. Model outputs are aggregated over multiple trials using a Bayesian framework to estimate the dog’s current likelihood of missing a target odor. A prior-informed thresholding step—adaptable to the training context, target odor, and prediction confidence—then supports decision-making: whether the dog should continue working or be given time to rest and recover.

To explore this question under controlled and reproducible conditions, we turned to the olfacto-treadmill system described earlier (10). This setup enables standardized exercise, precisely timed odor presentations, and high-quality video and physiological recordings—making it ideally suited for studying subtle pre-indication signals.

Importantly, we focused our analysis on the 4.5 seconds immediately preceding each target odor presentation. By restricting the input window to this pre-stimulus phase, we ensured that the dog’s behavior reflected its internal state rather than a direct response to the odor—something that is difficult to achieve in free-search environments.

Rather than analyzing raw video footage, we used markerless pose estimation to extract the trajectories of 29 anatomically defined joints and body landmarks (11). To improve the reliability of these pose predictions, we developed a segmentation-based preprocessing step that removes the background and focuses the algorithm’s attention on the dog. This enhancement significantly improved annotation quality—particularly for head and tail markers—and is critical for downstream modeling. While this adds some computational cost, the pipeline runs above 30 frames per second on standard hardware using MediaPipe (31), suggesting that real-time application is technically feasible and may be of interest to other researchers relying on pose-based analysis.

A central challenge in modeling detection performance lies in capturing the complexity of movement—not just what the dog is doing, but how it is moving, and how this evolves over time. Our predictive task relied solely on a brief 4.5-second segment—135 frames—recorded before any odor presentation, making it essential to extract the subtle spatial and temporal structure embedded in these sequences. To achieve this, we leveraged the fact that the tracked body points form a coherent skeletal structure: joints are not isolated measurements but part of a connected whole. This justified the use of a graph-based representation, where joints are treated as nodes and anatomical connections as edges. By applying a Graph Attention Network (GAT), we allowed the model to dynamically weigh the influence of each joint’s neighbors, capturing coordinated movement patterns that might signal fatigue or disengagement (33). Stacking two GAT layers further enabled the model to integrate information from more distal parts of the body, which may be important for recognizing whole-body posture shifts.

In parallel, we used a Temporal Convolutional Network (TCN) to capture how these postures change across time (34). This approach avoids the limitations of recurrent models while allowing the network to detect both short bursts of movement and longer behavioral trends. Together, the spatial and temporal modules enabled the model to extract fine-grained dynamics from anticipatory behavior—an essential capability when predicting performance before any overt alert behavior has occurred.

### Video and Physiology in Action: How Well the Model Performs and What It Sees

Having established a model capable of capturing structured movement dynamics, we next evaluated how well it could distinguish between trials that would result in an indication versus a missed alert. Despite relying solely on pre-stimulus movement data—with no access to odor cues or physiological signals—the video-only model achieved substantial predictive performance. It was particularly effective in identifying trials where the dog failed to respond: in 82% of cases where the model predicted a non-indication, it was correct, and it successfully detected 77% of all actual non-indications.

These results suggest that subtle changes in movement—occurring before the dog has encountered the odor—can already signal reduced readiness or engagement. However, movement alone only provides part of the picture. To probe internal state more directly, we extended the model to include physiological signals: heart rate, inter-beat interval, and core body temperature. This multimodal approach yielded further gains in predictive performance. Most notably, it improved the model’s ability to detect non-indication trials, correctly flagging 85% of all missed alerts while maintaining a high precision of 81%. This indicates that integrating autonomic markers with behavioral dynamics provides a more comprehensive and reliable assessment of the dog’s performance state.

While predictive accuracy is essential, interpretability is equally important—both to support handler trust and to guide future sensor development. This is particularly true for deep learning models, which are often criticized for their opaque decision processes (28). To address this, we applied two complementary strategies aimed at uncovering which inputs the model relied on most: saliency analysis for the pose data and ablation testing for the physiological signals. The saliency analysis perturbed individual joints to estimate how strongly each body point influenced the model’s output (35). Remarkably, the network focused most heavily on the head and tail regions—precisely the areas experienced handlers often monitor when assessing a dog’s motivation or engagement. That the model arrived at this focus without any explicit anatomical guidance suggests that these features emerged naturally from the data as informative indicators.

For the physiological inputs, we simulated sensor dropout by masking each signal independently. This revealed that heart rate variability, captured via inter-beat interval, had the strongest influence on the model’s decisions. In contrast, heart rate and core temperature—while commonly used as indicators of physical exertion—played a lesser role. This pattern may reflect that missed indications are not simply a matter of physical fatigue, but rather involve changes in arousal, stress, or cognitive load (23; 24). As such, IBI may serve as a sensitive marker of emerging mental fatigue or disengagement—internal states that precede overt behavioral decline.

### From Prediction to Decision Support: Tracking Performance with Bayesian Confidence

The interpretability analysis confirmed that the model’s predictions are not arbitrary: they rely on behaviorally and physiologically meaningful cues. Still, knowing whether a dog is likely to miss a specific indication is only part of the picture. From a handler’s perspective, it may be even more important to recognize when the dog’s overall performance begins to degrade—when it crosses a threshold beyond which continued deployment becomes risky. This shifts the focus from predicting isolated trial outcomes to estimating the dog’s internal state over time. Formally, this involves inferring a latent variable—such as the dog’s current miss rate—from a sequence of model outputs. A Bayesian framework provides a natural solution: by aggregating trial-level predictions, we can estimate a posterior distribution over the miss rate and quantify the probability that it exceeds an operationally relevant threshold. While the limited size of our dataset precluded full probability calibration, we nonetheless demonstrate how Bayesian updating supports interpretable, real-time risk estimation. Crucially, the framework is highly adaptable. The acceptable miss rate and the required confidence threshold can be tuned to match the operational context—more conservative for explosives detection in military scenarios, more flexible for agricultural or customs inspections. Prior knowledge can also be incorporated: the dog’s training history can inform the strength of the prior, while task-specific difficulty (e.g., target odor concentration) can shape the expected baseline performance. This makes the approach a practical and principled bridge between statistical modeling and deployment decisions, supporting handlers in making informed, context-sensitive choices in the field.

While our results demonstrate the potential of combining behavioral and physiological signals to detect early signs of performance decline, several limitations must be acknowledged. All data were collected under highly controlled laboratory conditions using the olfacto-treadmill—a valuable testbed that enabled precise measurement and standardization, but one that does not reflect the complexity of real-world deployments (10). In particular, the reliance on video-based pose estimation presents a practical challenge: continuous, unobstructed video capture is rarely feasible during operational searches. However, our findings suggest that the predictive signal lies in the movement pattern itself—not in the video modality per se. This opens the door to more practical sensing approaches, such as accelerometers, which are already well established in canine research and offer a robust, field-compatible alternative (16; 17; 18). A logical next step would be to replicate our analyses using such wearable sensor data, recognizing that this would require adapting the model architecture to accommodate different input characteristics.

Here, the olfacto-treadmill again proves its value: it provides a controlled environment for developing and validating sensor-based models before transitioning to more complex, semi-naturalistic settings. If consistent patterns can be captured from wearable data—and potentially enriched with physiological inputs—this would pave the way toward wearable systems capable of real-time readiness assessment. In this sense, our study lays the groundwork for practical, sensor-driven tools that support dog handlers in making informed, welfare-conscious deployment decisions.

## Supporting information

**S1 Video Segmentation-enhanced pose estimation.** The video shows a detection dog on a treadmill undergoing markerless pose prediction. The first segment displays the raw video. A semi-transparent background is then overlaid with joint predictions. In the final segment, the background is fully removed, illustrating the segmented condition used to improve pose estimation confidence and precision.

**S2 Video Visualization of keypoint saliency prior to odor presentation.** The video shows the output of markerless pose prediction for a detection dog on a treadmill. Each joint is color-coded to reflect its saliency—i.e., its influence on the model’s prediction of whether the dog will indicate. Saliency is shown dynamically across the 4.5-second window preceding odor presentation, with red indicating highly informative keypoints and blue indicating low saliency. The animation highlights how the model shifts attention between body regions over time.

**S1 Fig Full training metrics for video-only model.** Shown are training and validation curves for the ST-GAT + TCN model trained on 2D joint trajectories extracted from video, without physiological inputs. Metrics include binary cross-entropy loss, accuracy, AUC, precision, and recall across 60 training epochs. Each curve corresponds to the best-performing training run based on peak validation AUC. While training performance improved steadily, validation metrics began to fluctuate beyond epoch 45, indicating potential onset of overfitting.

**S2 Fig Full training metrics for multimodal model (video + physiology).** Shown are training and validation curves for the ST-GAT + TCN model trained on 2D joint trajectories combined with physiological inputs (heart rate, inter-beat interval, and core body temperature). Metrics include binary cross-entropy loss, accuracy, AUC, precision, and recall across 90 training epochs. Training and validation curves remained well aligned throughout, indicating stable learning and improved generalization compared to the video-only model.

**S1 Table Wilcoxon U test results comparing likelihoods of keypoint predictions between the full and segmented pipelines on the test set.** Asterisks indicate significant improvements (*p <* 0.05) in the segmented condition.

## Data and Code Availability

All code, trained models, and example data required to reproduce the analyses are available at Zenodo: 10.5281/zenodo.17197670 (version v1.0.0).

## Author Contributions

**Conceptualization:** Jörg Schultz, Edgar Aviles-Rosa, Nathaniel J. Hall, Michele N. Maughan

**Data curation:** Jörg Schultz

**Formal analysis:** Jörg Schultz

**Funding acquisition:** Michele N. Maughan

**Investigation:** Liza Rothkoff, Edgar Aviles-Rosa, Nathaniel J. Hall

**Methodology:** Jörg Schultz, Liza Rothkoff, Edgar Aviles-Rosa, Nathaniel J. Hall, Michele N. Maughan

**Project administration:** Michele N. Maughan

**Software:** Jörg Schultz

**Visualization:** Jörg Schultz

**Writing – original draft:** Jörg Schultz

**Writing – review & editing:** Jörg Schultz, Liza Rothkoff, Edgar Aviles-Rosa, Nathaniel J. Hall, Michele N. Maughan

## Acknowledgements

Funding was provided by the U.S. Army via the Chemical Biological Advanced Materials and Manufacturing Science (CBAMMS) Program (PE 0601102A Project VR9) at the Combat Capabilities Development Command Chemical Biological Center (DEVCOM CBC).

