## Supplemental Table 1 for "Detection dog performance state estimation from pre-stimulus video and physiological signals using deep learning and Bayesian inference"

### S1 Table. Wilcoxon U test results for per-keypoint likelihoods.

| ID | Keypoint | U Statistic | p-value | Direction / Significance |
|---|---|---|---|---|
| 1 | BodyPart1 | 5 | 3e-05 | full < segmented* |
| 2 | BodyPart2 | 0 | 1e-05 | full < segmented* |
| 3 | BodyPart3 | 2 | 1e-05 | full < segmented* |
| 4 | BodyPart4 | 51 | 0.04529 | full < segmented* |
| 5 | BodyPart5 | 115 | 0.06197 | full > segmented |
| 6 | BodyPart6 | 125 | 0.02012 | full > segmented* |
| 7 | BodyPart7 | 74 | 0.30404 | full < segmented |
| 8 | BodyPart8 | 72 | 0.26915 | full < segmented |
| 9 | BodyPart9 | 60 | 0.1092 | full < segmented |
| 10 | BodyPart10 | 20 | 0.00052 | full < segmented* |
| 11 | BodyPart11 | 28 | 0.00204 | full < segmented* |
| 12 | BodyPart12 | 18 | 0.00036 | full < segmented* |
| 13 | BodyPart13 | 57 | 0.08309 | full < segmented |
| 14 | BodyPart14 | 55 | 0.06848 | full < segmented |
| 15 | BodyPart15 | 20 | 0.00052 | full < segmented* |
| 16 | BodyPart16 | 12 | 0.00011 | full < segmented* |
| 17 | BodyPart17 | 50 | 0.04062 | full < segmented* |
| 18 | BodyPart18 | 80 | 0.60118 | full > segmented |
| 19 | BodyPart19 | 87 | 0.45915 | full > segmented |
| 20 | BodyPart20 | 3 | 2e-05 | full < segmented* |
| 21 | BodyPart21 | 24 | 0.00105 | full < segmented* |
| 22 | BodyPart22 | 39 | 0.01051 | full < segmented* |

| ID | Keypoint | U Statistic | p-value | Direction / Significance |
|---|---|---|---|---|
| 23 | BodyPart23 | 50 | 0.04062 | full < segmented* |
| 24 | BodyPart24 | 91 | 0.37916 | full > segmented |
| 25 | BodyPart25 | 29 | 0.0024 | full < segmented* |
| 26 | BodyPart26 | 33 | 0.00446 | full < segmented* |
| 27 | BodyPart27 | 50 | 0.04062 | full < segmented* |
| 28 | BodyPart28 | 38 | 0.00916 | full < segmented* |
| 29 | BodyPart29 | 156 | 0.00014 | full > segmented* |

**Statistically significant at $p < 0.05$*
