## Supplementary figures and images for "Detection dog performance state estimation from pre-stimulus video and physiological signals using deep learning and Bayesian inference"

### Supplemental Figure 1

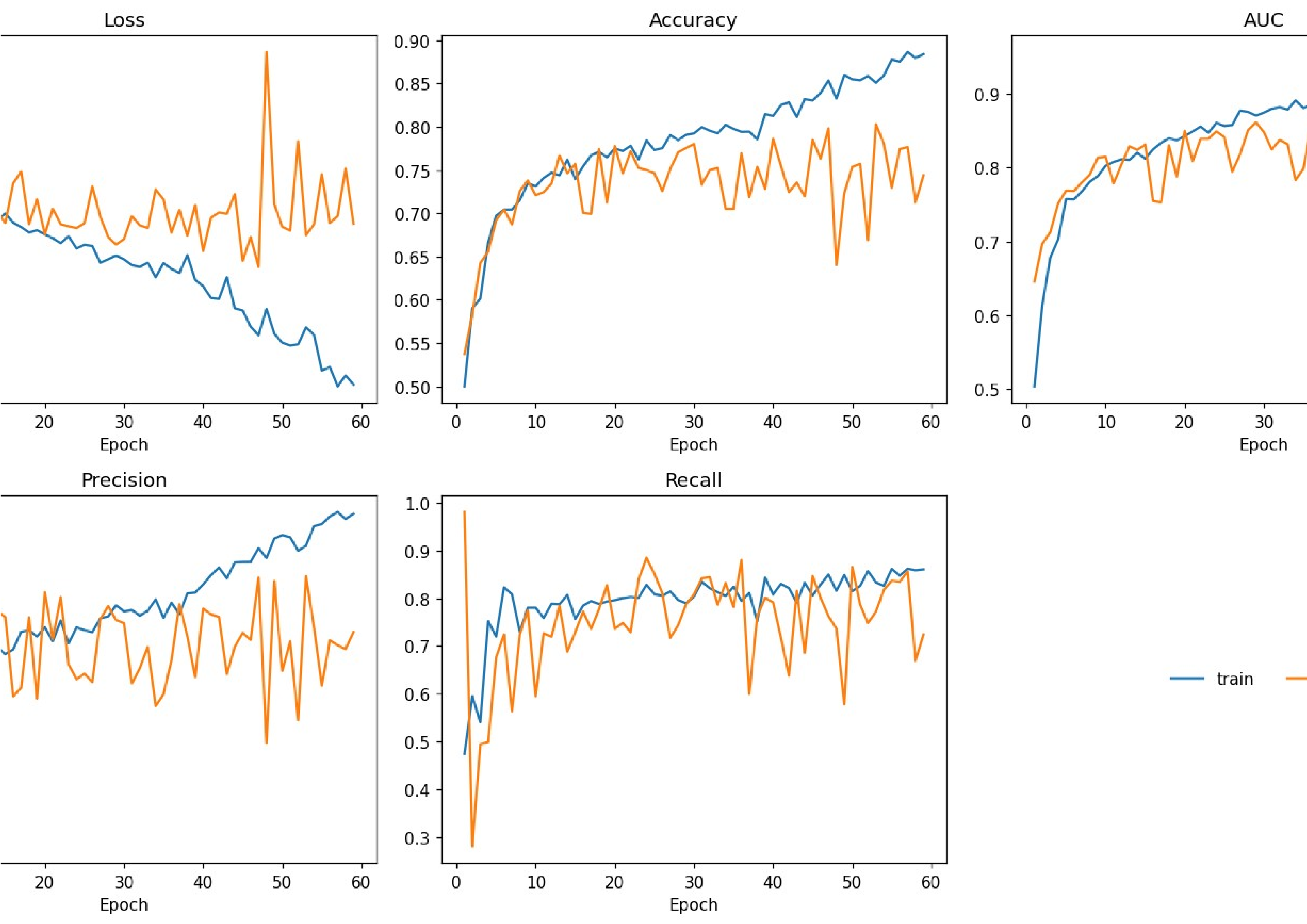

### Supplemental Figure 2

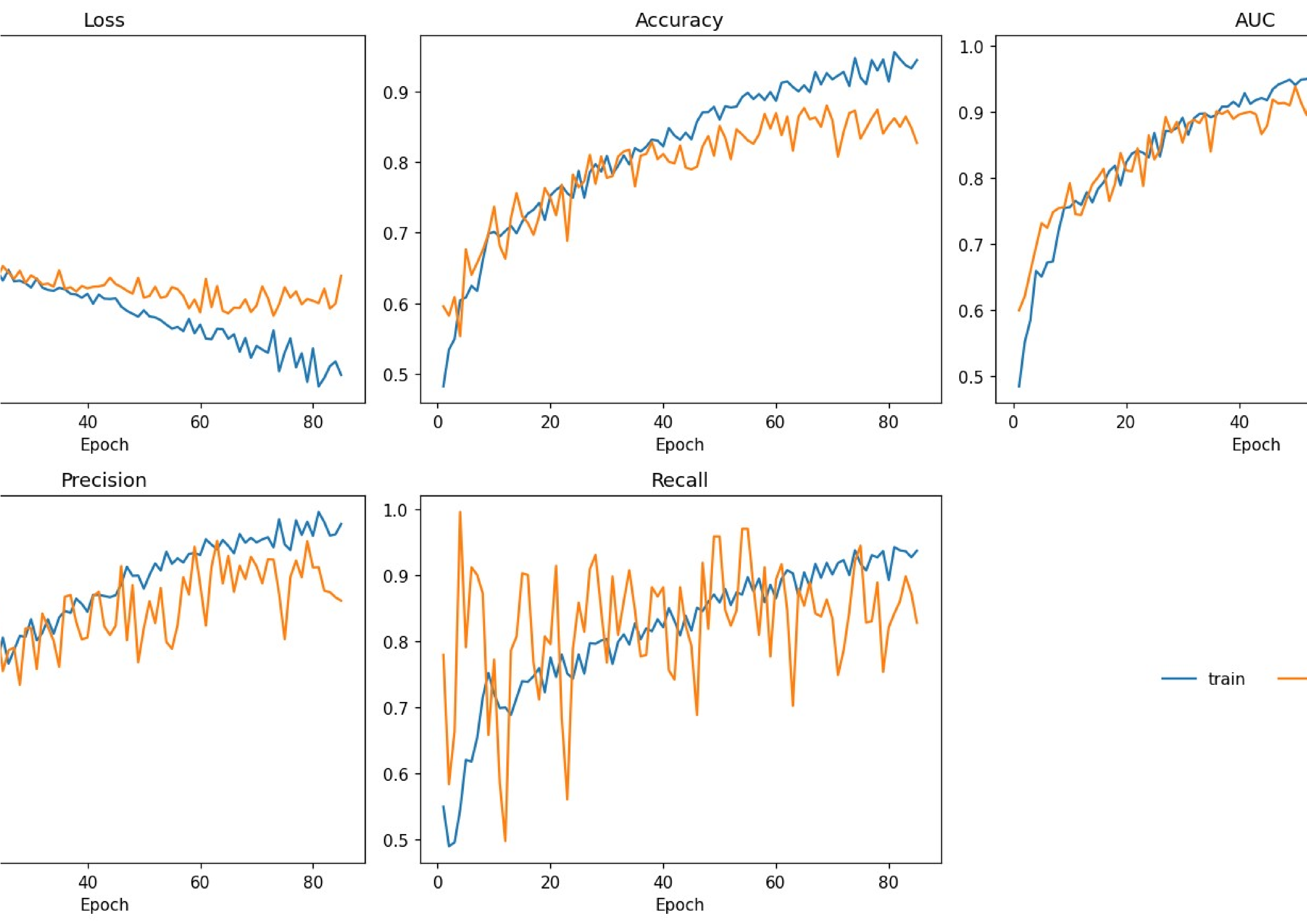
